# A mechanical circuit in End4p coordinates force transmission during clathrin-mediated endocytosis

**DOI:** 10.1101/2023.10.23.563344

**Authors:** Yuan Ren, Jie Yang, Barbara Fujita, Yongli Zhang, Julien Berro

## Abstract

Mechanical forces are transmitted from the actin cytoskeleton to the membrane during clathrin-mediated endocytosis (CME) in the fission yeast *Schizosaccharomyces pombe*. The onset and termination of force transmission is tightly regulated temporally during different stages of CME, and spatially over the surface of the invaginated membrane. How force transmission is regulated and coordinated at the molecular scale is unclear. An adaptor protein in CME, End4p, directly transmits force by binding to both the membrane (through ANTH domain) and F-actin (through THATCH domain). We show that 8pN is required for stable binding between THATCH and F-actin. We also report the discovery and characterization of a new domain on End4p, which we named Rend (<u>R</u> domain in <u>End</u>4p), that resembles R12 of talin. Membrane localization of Rend primes the binding of THATCH to F-actin, and force-induced unfolding of Rend at 15pN terminates the transmission of force during CME. We show that the mechanical properties (mechanical stability, unfolding length, hysteresis) of Rend and THATCH are tuned to form an auto-regulated circuit for the initiation, transmission and termination of force between the actin cytoskeleton and membrane. Shorting the circuit leads to permanent End4p association with the membrane or with F-actin, or failure to enter the force transmission cycle. Mathematical modeling of force transmission through Rend-THATCH connection shows that input force from F-actin is buffered to a narrow range towards the membrane. The mechanical circuit by Rend and THATCH may be conserved and coopted evolutionarily in cell adhesion complexes.

## Background

Mechanical forces cannot be transmitted without physical connections. For cells, connections between cell membrane and the actin cytoskeleton provides the linkage for the transmission of force in cell-cell adhesion, cell-matrix attachment, and for the internalization of cell membrane during endocytosis(*1–4*). Force-bearing components of these mechanical connections include cadherin/catenin, fibronectin/integrin/vinculin, and epsin/HIP1R, among others(*5–9*). Dozens of regulatory proteins fine tune the composition and the strength of mechanical connections in response to both chemical (e.g., ligand type and density) and mechanical cues (e.g., substrate rigidity, membrane tension)(*10–14*), and faulty connections are implicated in a variety of developmental and physiological disorders(*1*, *15*, *16*).

The need to construct stable and adaptive supramolecular mechanical connections poses a challenge for adhesion protein machineries. Proteins have to bind tightly to sustain force transmission, but may also need to unbind quickly to terminate force transmission to meet dynamic mechanical needs, i.e., produce the right amount of force at the right place at the right time. Growing data suggest that mechanical forces alter the binding between proteins in non-intuitive ways(*2*, *17*, *18*). For example, several of the F-actin binding proteins in cell adhesion (α-catenin, talin, vinculin) have been found to display catch bond behavior, where force increases their binding to F-actin, especially when pulled towards the pointed end of F-actin(*19–22*). Moreover, domains in talin and α-catenin can be mechanically unfolded to reveal cryptic binding motifs that are not accessible to binding partners when folded but, when unfolded, can selectively recruit proteins such as vinculin to strengthen focal adhesions(*23–27*). Interestingly, the F-actin binding domains in cell adhesion proteins are often five-helix-bundles(*18*, *28*), and force induced partial unfolding of the five-helix-bundle has been recently shown to underlie the catch bond mechanism(*26*). Deletion of the first alpha-helix from αE-catenin eliminated the catch bond behavior, resulting in stable binding to F-actin that can be described by a single state slip bond(*26*). Whether this is a conserved mechanism to form catch bonds in other mechanotransduction pathways is unknown, and *in vivo* data to compare the force transmission with or without catch bond formation are missing. In addition, the catch bond mechanism from F-actin binding domains alone is insufficient to explain how the initial binding of these adhesion proteins is achieved in the absence of force when the binding affinity is low, and what stops the high affinity binding after stable catch bonds are formed.

End4p is a homolog of HIP1R in the fission yeast *Schizosaccharomyces pombe*, and the F-actin binding domain in End4p is a five-helix-bundle named THATCH (**t**alin-**H**IP1/R/Sla2p **a**ctin-**t**ethering **C**-terminal **h**omology)(*28–32*). End4p transmits mechanical forces from F-actin to the plasma membrane during clathrin-mediated endocytosis (CME), and we have recently shown that there is a gradient of force along End4p molecule during CME *in vivo*(*7*, *33*). The peak force before THATCH reaches ∼19pN, which drops to ∼11pN near the clathrin lattice and ∼9pN near the membrane(*7*). Compared with cell-cell and cell-matrix adhesion that exists in the minutes to hours’ time scale, CME in the fission yeast is transient, where F-actin driven formation of a vesicle out of an initially flat membrane only lasts ∼10 seconds for each endocytic event(*3*, *4*). Therefore, the binding between F-actin and End4p must be tightly coupled spatially and temporally to ensure robust onset and termination of force transmission. How is this achieved is unclear. Here, we show that the physical connection between two mechanosensitive domains, Rend and THATCH, acts as a mechanical circuit to start force transmission only at the plasma membrane, to buffer the transmission of force to a small range, and to terminate force transmission at a hard-wired magnitude. This circuit may represent a general solution to regulate catch bonds in force transmission.

## Results

End4p forms a dimer in fission yeast cells(*33*, *34*). End4p binds to the plasma membrane via its N-terminal ANTH (**A**P180 **N**-**t**erminal **h**omology) domain, and to F-actin through the C-terminal THATCH domain(*29*, *35–37*). End4p interacts with the clathrin lattice by binding to clathrin light chain and binds to several SH3 domain-containing proteins through its proline rich domain (PRD) (Fig. 1a)(*34*, *38–40*). The extensive interactions between End4p and other endocytic proteins relay forces between F-actin and the plasma membrane(*7*).

**Figure 1.**
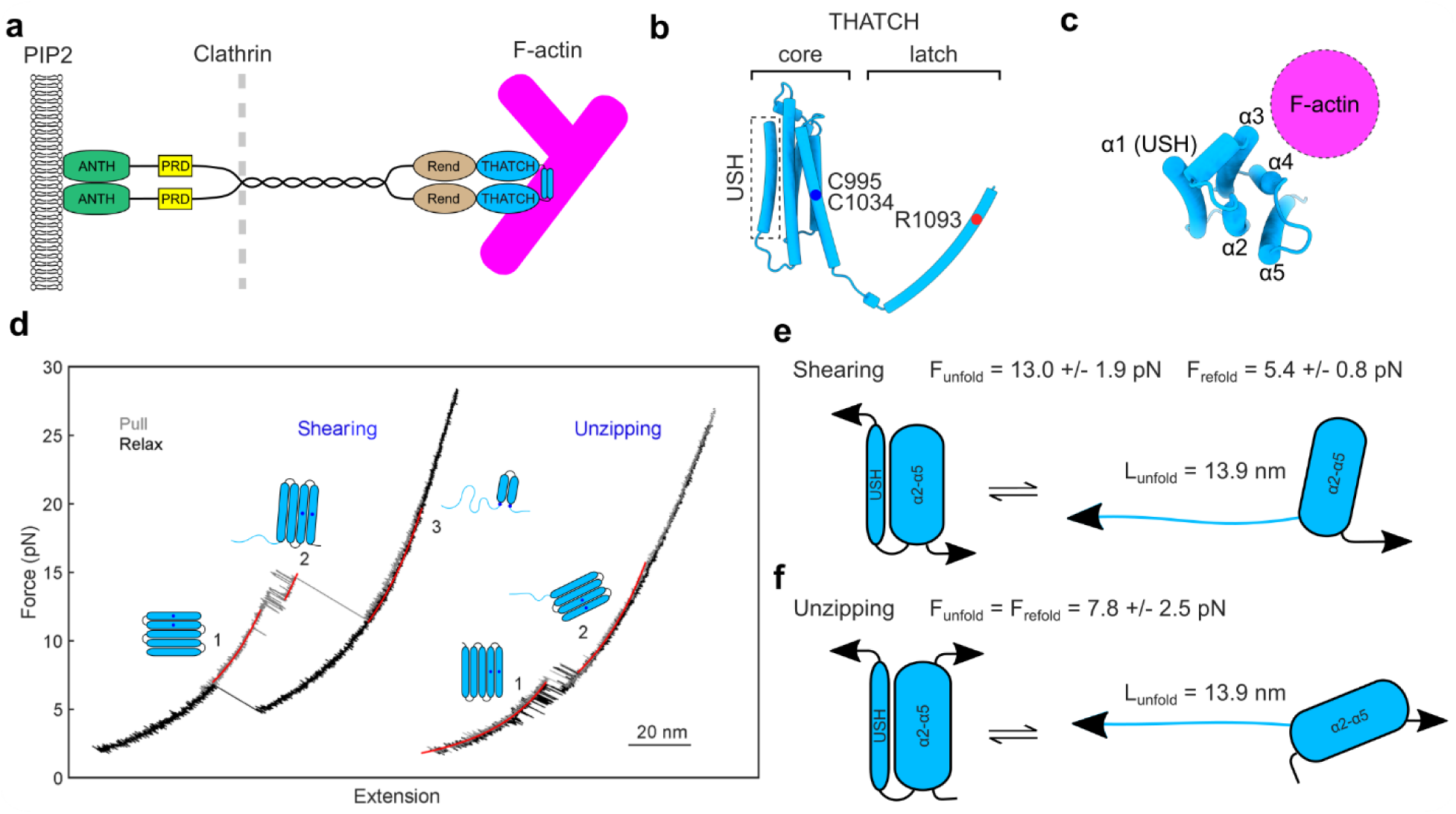
The unfolding of USH from THATCH requires at least 8pN force. **a**, Schematic of an End4p dimer between the lipid membrane and the actin cytoskeleton. The N-terminal ANTH domain binds to PIP2. The proline-rich domain (PRD) binds to SH3 domain-containing protein in the endocytic coat. An unknown motif before the dimerization domain binds to clathrin light chain. THACH domain binds to F-actin. Drawing is not to scale. **b**, Structure of THATCH from End4p as predicted by AlphaFold. THATCH is made from a five-helix-bundle (core) and a C-terminal tail (latch) that mediates dimerization. The first alpha-helix of THATCH core is termed USH. An arginine (R1093) on latch is critical for dimerization. Two cysteine (C995 and C1034) residues form a disulfide bond. **c**, The F-actin-binding surface of THATCH core is located at the opposite side of USH based on previous structural data(*28*, *32*). The barbed end of F-actin is facing the reader in this configuration. Note that pulling USH towards the pointed end of F-actin, or away from F-actin, leads to unzipping of USH. **d**, Force-extension curves (FEC) of THATCH core pulled in the shearing or the unzipping orientation. Different states of THATCH core are placed next to the corresponding curve. See full analysis of FEC in Fig. S2. **e**, USH unfolds at 13 ± 1.9pN (mean ± SD) and refolds at 5.4 ± 0.8pN when pulled in the shearing orientation. **f**, USH unfolds and refolds at an equilibrium force of 7.8 ± 2.5pN when pulled in the unzipping orientation, which reflects the F-actin binding orientation in **c**.

The THATCH domain is composed of a “core” that directly binds to F-actin and a “latch” that mediates dimerization by forming an anti-parallel coiled-coil (Fig. 1b; Fig. S1)(*28*, *32*). The structures of THATCH from human HIP1R and mouse talin-1 reveal a F-actin binding surface at the opposite side of the first alpha-helix (also named upstream helix or USH) (Fig. 1c). Dimerization through latch greatly increases the binding affinity between THATCH core and F-actin, and the presence of USH strongly inhibits binding, as shown from bulk biochemical assays(*28*, *32*, *41*, *42*). Dimerized THATCH from human talin-1 forms an asymmetric catch bond with F-actin, where force >5pN on THATCH towards the pointed end of F-actin is required for stable binding(*22*).

We began by measuring the mechanical stability of the purified THATCH core from End4p using optical tweezers. Unlike previous experiments where the binding between F-actin and THATCH is studied in reconstituted system, we focused solely on the response of isolated THATCH domain to tensile forces. The two termini of the THATCH core were connected to two optically trapped beads, and force on the THATCH core was changed by controlling the distance between the two beads. When the THATCH core was pulled from its original N- and C-termini, it partially unfolded when the force reached 13 ± 1.9pN (mean ± SD) and refolded when force dropped below 5.4 ± 0.8pN (mean ± SD) (Fig. 1d-e). During THATCH core unfolding, we first observed a transient state (state 2 in Fig 1d), which corresponded to the unfolding of USH, before we observed a transition to a state where all but two helices maintained together by a disulfide bond unfolded (state 3 in Fig 1d, Fig. S2). By introducing a cysteine into the loop between the 4^th^ and 5^th^ alpha-helices to form a disulfide bond with the bead linker, we were able to mimic the force applied by F-actin on the THATCH core. In this configuration, the THATCH core unfolded and refolded at an equilibrium force of 7.8 ± 2.5pN (mean ± SD), and the unfolding distance matched the contour length of USH (Fig. 1d, 1f; Fig. S2). No further unfolding event was observed up to ∼25pN (Fig. 1d) showing the remaining 4 helices of the THATCH core can sustain large forces without unfolding. The unfolding of USH is reversible in both the shearing and the unzipping mode, suggesting fast transitions between the folded and the unfolded states of the THATCH core under tension. Altogether, our data suggest that a THATCH core where USH is mechanically dissociated from the remaining four-helix-bundle (state 2), can be achieved either by forces applied on the C-terminus of the THATCH domain or from the F-actin binding surface, and at least ∼8pN force is needed to maintain the THATCH core in an open conformation.

To explore the biological function of the USH domain, we went on to study the affinity of THATCH constructs to F-actin with or without USH in fission yeast cells. In wild-type cells, full-length End4p localizes at endocytic sites whose location correlated with sites of cell wall deposition, i.e., at the cell tips during cell growth and at the midline during cell division (Fig. 2a)(*7*, *43*). Isolated THATCH dimers displayed a diffusive localization throughout the fission yeast cell, indicating a low affinity to F-actin, similarly to THATCH homologues (Fig. 2b)(*44*, *45*). In contrast, overexpression of THATCH-ΔUSH dimers induced the formation of thick F-actin bundles and large puncta (Fig. 2c)(*34*). Both structures contained actin crosslinker fimbrin (Fim1p) but not alpha-actinin (Ain1p) (Fig. 2d-e, movie 1). The presence of filament bundles depended on THATCH-ΔUSH dimerization, as monomeric constructs which contained a point mutation preventing dimerization of the latch (THATCH-ΔUSH-R1093G) were diffusive (Fig. 2f). The F-actin structures induced by THATCH-ΔUSH dimers were remarkably stable, as treatment with Latrunculin A at a dose that destroys all F-actin structures in wild-type fission yeast cells(*46*) (Fig. 2g) had negligible effect on the F-actin bundles and puncta. To test the activity of THATCH-ΔUSH dimers at the native expression level, we used the 2A peptide to create strains where THATCH-ΔUSH dimers and the rest of End4p (containing or not containing the USH domain) were expressed in the same cell with equal stoichiometry. F-actin bundles and puncta only formed in cells with End4 constructs that did not contain USH (Fig. 2h-i). These data demonstrate that USH is a potent inhibitor of the F-actin-binding ability of THATCH-ΔUSH dimer, and that constitutively active THATCH, even at the native expression level, creates overly stable F-actin structures in fission yeast cells. Combining these results with data from *in vitro* force spectroscopy (Fig 1), our data suggest that force >8pN turns THATCH from an “OFF” state where its affinity to F-actin is negligible, to an “ON” state where THATCH strongly bundles F-actin (Fig. 2j). Therefore, mechanical forces on THATCH directly regulates its affinity to F-actin by controlling the unfolding/refolding of USH.

**Figure 2.**
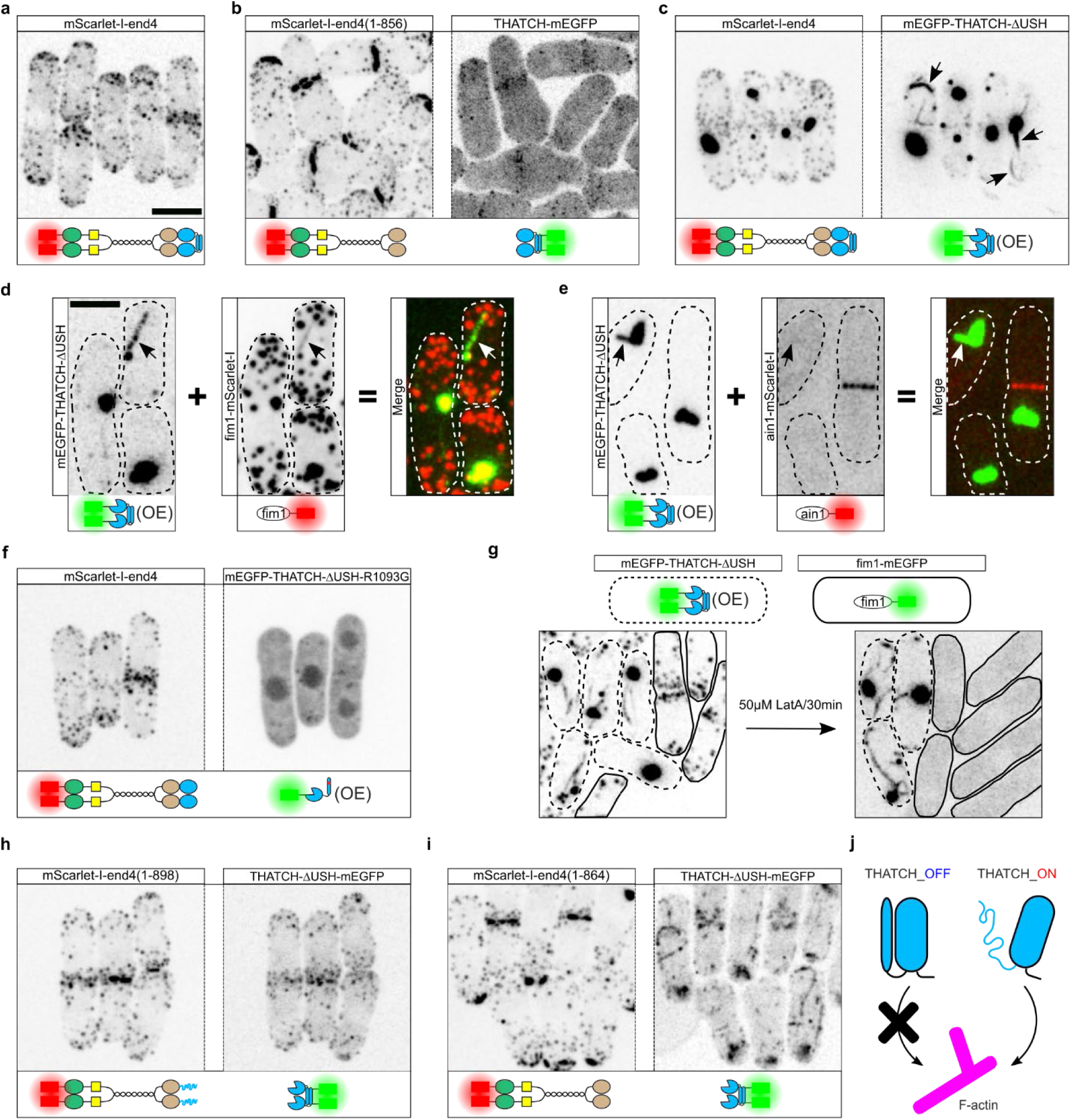
USH inhibits the binding of THATCH to F-actin *in vivo*. **a**, mScarlet-I tagged End4p localizes to endocytic sites in fission yeast cells. Endocytic sites are enriched at the cell tips and the division plane. **b**, THATCH is disconnected from End4p by a self-cleaving 2A peptide, and fragments of End4p are visualized by two different fluorescent tags. The N-term fragment (tagged by mScarlet-I) forms puncta at cell tips and the division plane, and isolated THATCH (tagged by mEGFP) is diffusive in the cytoplasm, indicating a lock of affinity to F-actin. **c**, mEGFP-THATCH-ΔUSH is overexpressed in fission yeast cells where wild type End4p is tagged with mScarlet-I. The overexpression of mEGFP-THATCH-ΔUSH caused the formation of large puncta and bundles (arrows). Wild type End4p is recruited to the puncta but not the bundles. See also movie 1. **d**, mEGFP-THATCH-ΔUSH is overexpressed in fission yeast cells where actin crosslinker Fim1p is tagged with mScarlet-I. Fim1p co-localizes with both the puncta and bundle. **e**, mEGFP-THATCH-ΔUSH is overexpressed in fission yeast cells where actin crosslinker Ain1p is tagged with mScarlet-I. Ain1p shows no co-localization with either the puncta or bundle. **f**, mEGFP-THATCH-ΔUSH-R1093G is overexpressed in fission yeast cells where wild type End4p is tagged with mScarlet-I. R1093G mutation prevents the dimerization of THATCH. Monomeric THATCH is diffusive in the cytoplasm without forming puncta or bundles. **g**, 50 μM Latrunculin A (LatA) was used to disassemble F-actin in two populations of fission yeast cells. Cells enclosed by dotted line has overexpression of mEGFP-THATCH-ΔUSH, and cells enclosed by solid line expresses Fim1p-mEGFP. Note that puncta and bundles are insensitive to LatA treatment, whereas Fim1p became diffusive in the cytoplasm. **h**, A self-cleaving 2A peptide was inserted after USH to disconnect THATCH-ΔUSH (tagged by mEGFP) from the N-term fragment of End4p (tagged by mScarlet-I). USH inhibited the formation of puncta and bundles by THATCH-ΔUSH even when it is physically disconnected from THATCH-ΔUSH. **i**, A self-cleaving 2A peptide was inserted after USH to disconnect THATCH-ΔUSH (tagged by mEGFP) from the N-term fragment of End4p (tagged by mScarlet-I), and USH is deleted from End4p. The absence of USH led to constitutively active THATCH-ΔUSH which drove the formation of puncta and bundles. **j**, THATCH exits in two states each with different affinities to F-actin. THATCH_OFF contains folded USH and has negligible affinity to F-actin, and THATCH_ON has unfolded USH and strongly binds F-actin. Note that THATCH is drawn as a monomer for simplicity, but dimerization of THATCH is needed for binding to F-actin. Images are maximum intensity projections of the whole fission yeast cells. The schematic under each image indicates the modifications on End4p. Proteins are expressed at the endogenous level unless when labeled OE (overexpression). Scale bar in **a** applies to all images except **d**-**e**, 5 μm. Scale bar in **d** applies in **d-e**, 4 μm.

The peak force on THATCH *in vivo* is ∼19pN and large enough to unfurl USH. To turn THATCH off, tension of USH must drop below 8pN. This cannot be achieved by unfolding the remaining four-helix-bundle, which is stable up to at least ∼25pN (Fig. 1d; Fig. S2), or by disassembling F-actin, because F-actin bound by active THATCH is resistant to disassembly (Fig. 2c, 2g, 2i). A plausible way to reduce tension is to have other domains that unfold under tension, so that force on THATCH could be buffered. In talin, 13 rod domains (also called R domains) have been found to unfold with varying stabilities in the 5-25pN range. The unfolding of R domains buffers tension on talin to 5-10pN, despite significant changes in the total length of talin or fluctuations in talin extension. While comparing End4p sequences, we found a structured domain located in front of THATCH (Fig. 1a; Fig. S1). A previous study noticed the similarity between this domain and talin THATCH(*46*). This domain is predicted to be a five-helix-bundle by both AlphaFold and RaptorX (Fig. 3a). We verified the all-helical content of this domain through circular dichroism (Fig. S3a). This domain is conserved in all species that contain End4p/Sla2/HIP1R at the structural level but not the sequence level (Fig. S3b-c), and the predicted structure is most similar to that of R12 from talin, which is also the R domain right before THATCH (Fig. 3a)(*27*, *47*). The five alpha-helices of Rend are not organized the same way as THATCH’s helices. While consecutive helices of THATCH contact each other, the first helix of Rend does not contact helix 2 but helices 3 and 4, making the first alpha-helix of Rend less isolated than THATCH’s first helix (USH) (Fig. 3b). Consistent with this observation, force on purified Rend led to all-or-none unfolding (unfolding length 59nm) when the pulling force reached 14.8 ± 1.2pN (mean ± SD), instead of partial unfolding. Rend refolds with a strong hysteresis when the force drops to ∼2pN (Fig. 3c-d; Fig. S4). The mechanical properties of Rend indicate that tension before THATCH can only briefly exceed 15pN before Rend ruptures, and that tension cannot increase again before decreasing to ∼2pN for Rend to refold. USH refolds when the tension decreases below 8pN, which turns THATCH off and leads to disassociation from F-actin (Fig. 3e). As a result, Rend limits the transmission of tension to 15pN and effectively turns THATCH off once this limit is reached. We calibrated a new coiled-coil force sensor, cc-14pN (Fig. S5), which showed that the peak force before Rend *in vivo* is between 14 and 18pN (Fig. 3f-h), in agreement with our force buffering model.

**Figure 3.**
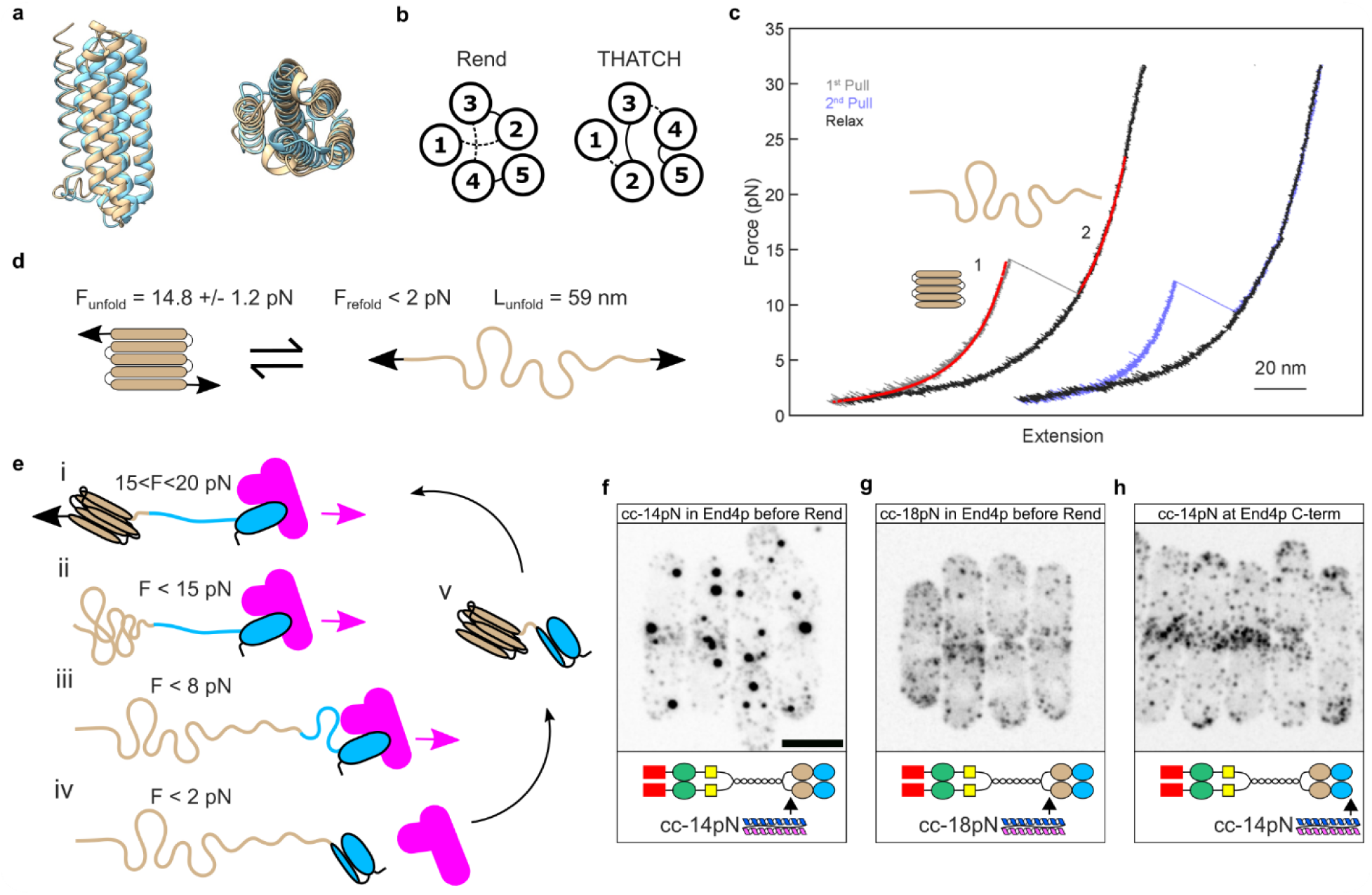
Rend unfolds at 15pN to terminate force transmission from THATCH. **a**, Predicted structure of Rend (mustard) docked to the crystal structure of R12 from talin (cyan, PDB:3dyj.), in two orthogonal views. Rend most resembles R12 among the 13 R domains of talin (RMSD: 2.2 Å). **b**, The order of connections of the five alpha-helices are different in Rend than in THATCH. Alpha-helices are represented as numbered circles. Solid lines indicate connections on top of alpha-helices and dotted lines indicate connections at the bottom. **c**, Representative FECs of Rend under tension. Two consecutive pulling events from the same molecule are shown to demonstrate the complete refolding of Rend after relax. See full analysis of Rend unfolding in Fig. S4. **d**, Rend unfolds at 14.8 ± 1.2pN (mean ± SD) and refolds at <2pN, with an unfolding distance of 59nm. Tensile force on protein domains directly connected to Rend is therefore buffered at 14.8pN, unless the full extension of unfolded Rend is achieved. **e**, Different states of Rend-THATCH domains are listed from top to bottom, and the tension between Rend and THATCH is indicated by F. i) forces from F-actin are transmitted through THATCH_ON to Rend until transiently reaching a peak magnitude 18-20pN(*7*); ii) force reaches the unfolding threshold of Rend, and Rend unfolds. Tension between Rend and THATCH_ON starts to drop; iii) the long unfolding distance of Rend (59nm), and the fast refolding of USH, prevents its full extension in CME, and tension between Rend and THATCH continues to drop. USH refolds when tension drops to below 8pN, and THATCH loses its binding to F-actin. iv) Rend refolds after tension is below 2pN; v) End4p with refolded Rend and THATCH could participate in force transmission cycle again. The mechanical unfolding of Rend inactivates THATCH and buffers force transmission before Rend. Pulling forces are indicated by arrows. **f**, A calibrated coiled-coil (cc-14pN) is inserted into End4p before Rend, and End4p is fluorescently tagged. The formation of End4p droplets indicates that force before Rend is large enough to unfold cc-14pN. **g**, A calibrated coiled-coil (cc-18pN) is inserted into End4p before Rend, and End4p is fluorescently tagged. The absence of End4p droplets indicates that peak force before Rend is below 18pN. **h**, cc-14pN is tagged to the C-terminus of fluorescently labeled End4p. cc-14pN did not drive the formation of End4p droplets at the absence of force. The schematic under each image indicates the modifications on End4p. Proteins are expressed at the endogenous level. Scale bar in **f** applies to **f**-**h**, 5 μm.

We also discovered that Rend has an additional function in organizing End4p molecules *in vivo*. End4p constructs that are disconnected from THATCH formed large membrane associated puncta in fission yeast cells, whereas End4p constructs disconnected from Rend-THATCH did not (Fig. 4a-b). End4p with the deletion of Rend (end4-ΔRend) was more cytoplasmic than wild type End4p (Fig. 2c-d, 2f) and has impaired localization to endocytic sites (Fig. 4g), suggesting that Rend increases the local concentration of End4p. Rend deletion also slowed cell growth (Fig. S6). We saw similar defects after replacing Rend with R3 from talin (Fig. 4e-g, S6). To test if Rend could cause protein puncta formation independent of other domains from End4p, we compared the localization of isolated Rend constructs in fission yeast cells. Overexpressed Rend dimer was diffusive in the cytoplasm (Fig. 4h). A membrane localization signal (single copy of the PH domain from fission yeast Plc1p) was sufficient to induce the formation of Rend puncta from dimerized but not monomeric Rend (Fig. 4i-j). In the same vein, attachment of dimerized but not monomeric Rend to the C-terminus of a BAR domain containing membrane protein, Pil1p, reshaped the linear furrows (eisosome) into spherical puncta (Fig. S7). These data demonstrate that Rend could mediate protein puncta formation at the plasma membrane in an autonomous fashion. *In vivo* protein condensation is often under phosphor-regulation. Since Rend contains a threonine (T841) that loosely conforms to the recognition motif of Ark1/Prk1 family kinase(*48–50*) (Fig. S8), we wondered if the phosphorylation state of T841 could control the formation of puncta *in vivo*. We tested this hypothesis by constructing a phospho-mimetic and a non-phosphorylatable Rend (respectively T841E and T841A). The phospho-mimetic Rend did not form puncta, while the non-phosphorylatable Rend did (Fig. 4k-l), demonstrating that phosphorylation state of T841 can control the assembly and disassembly of Rend and End4p. The ability to form protein puncta is not a general property of all R domains, as only R12 but not R3 from talin formed protein puncta when dimerized and attached to PH domain (Fig. 4m-n). Taken together, our data demonstrate that membrane localization of Rend leads to strong self-association of Rend that increases its local concentration, possibly through condensation (Fig. 4o). Consequently, in End4p, Rend promotes the activation of THATCH in at least two ways: 1) increased concentration of THATCH leads to higher probabilities of binding to F-actin even at the low affinity OFF state; 2) because the motion of Rend is restricted after self-association, force from activated THATCH will be channeled through the dimerization tail to activate THATCH of other End4p molecules (Fig. 4o). By linking THATCH to Rend, THATCH is only activated at the endocytic site when a sufficient amount of End4p molecules is recruited, and Rend spatially coordinates the activation of THATCH.

**Figure 4.**
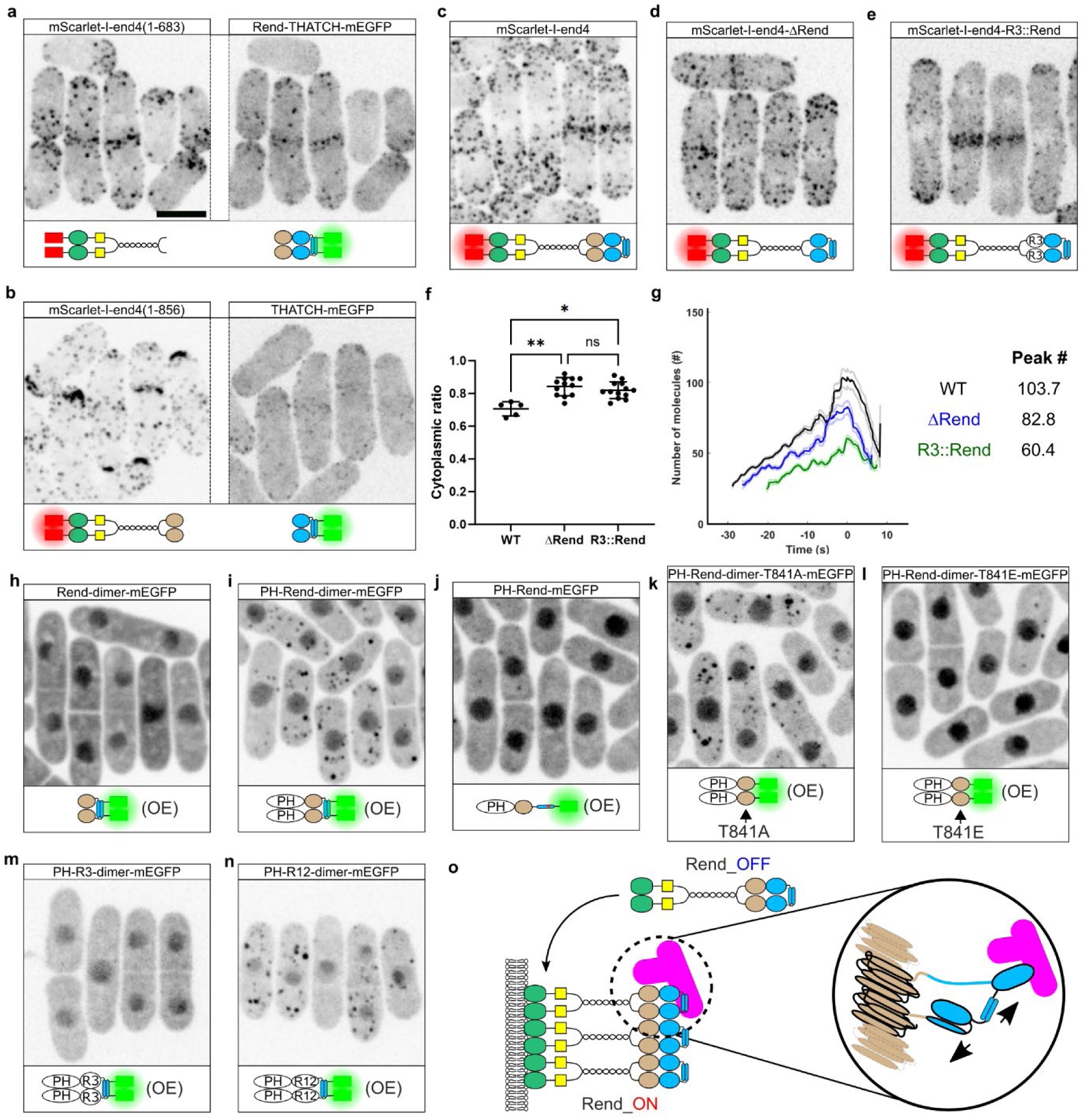
Membrane proximity drives the formation of Rend puncta. **a-b**, Rend-THATCH (**a**) or THATCH (**b**) is disconnected from End4p by a self-cleaving 2A peptide, and fragments of End4p are visualized by two different fluorescent tags. The N-term fragment (tagged by mScarlet-I) forms puncta at cell tips and the division plane only when it contains Rend, and isolated Rend-THATCH (tagged by mEGFP) is diffusive in the cytoplasm. **c**, Wild type End4p with a fluorescent tag. **d**, Rend is deleted from fluorescently tagged End4p. End4p-ΔRend has increased partition in the cytoplasm. **e**, Rend is replaced by R3 from talin, and End4p is fluorescently tagged. Replacement of Rend by talin R3 increased the partition of End4p in the cytoplasm. **f**, Quantification of the cytoplasmic ratio of End4p for **c-e**. The deletion of Rend increased End4p cytoplasmic ratio, and talin R3 could not rescue this phenotype. Kruskal-Wallis test. *: p<0.05; **: p<0.005. **g**, Temporal evolution of the number of End4p molecules at endocytic patches as determined by patch tracking. The deletion of Rend, or the replacement of Rend by talin R3, inhibited the accumulation of End4p molecules at endocytic sites. **h**, Overexpressed Rend dimer is diffusive in fission yeast cytoplasm. Rend dimerization is mediated by the latch from End4p THATCH domain. **i**, Overexpressed Rend dimer with a membrane localizing signal (PH domain) led to the formation of droplets in the cytoplasm. **j**, Overexpressed monomeric PH-Rend is diffusive in the cytoplasm and did not form droplets. The R1093G mutation in latch from End4p THATCH prevents dimerization. **k**, T841A phospho-dead mutation on overexpressed PH-Rend had no influence on the formation of droplets. **l**, T841E phosphor-mimetic mutation on overexpressed PH-Rend inhibited the formation of droplets. **m**, Overexpressed PH-R3 dimer did not form droplets in fission yeast cells. **n**, Overexpressed PH-R12 dimer formed droplets in fission yeast cells. **o**, Rend_OFF is converted to Rend_ON by localization to the membrane. Membrane-induced formation of Rend puncta primes the binding between THATCH and F-actin in two ways: 1) locally increasing the concentration of THATCH; 2) restricting the movement of THATCH, so that force from stochastically activated THATCH can be used to activate nearby THATCH. Scale bar in **a** applies to all yeast cell images, 5 μm.

We propose that two domains of End4p, Rend and THATCH, form a mechanical circuit that regulates the transmission of force from F-actin to the plasma membrane during endocytosis (Fig. 5a). Both Rend and THATCH have an ON and an OFF state (Rend_ON, Rend_OFF, THATCH_ON and THATCH_OFF). Rend is turned ON when it is folded and makes puncta at the membrane, and turned OFF when it is unfolded by >15pN force. THATCH is turned ON when the >8pN force unfolds USH, increasing its affinity to actin, and turned OFF when force drops <8pN and USH refolds with the rest of THATCH into a 5-helix bundle. A cycle of force transmission involves the sequential changes in the ON and OFF states of Rend and THATCH, where the magnitude of force triggers the transition between states (Fig. 5b). We tested this model by artificially changing the ON and OFF states of Rend and THATCH. Our model predicts that THATCH deletion results in an End4p with a constitutive THATCH_OFF, unable to apply force on Rend, therefore unable to remove Rend from the membrane. This was confirmed by experiments showing large aggregates in end4-ΔTHATCH mutants (Fig. 5c). Force on Rend can also be removed by preventing the dimerization of THATCH or by preventing actin assembly with Latrunculin A (LatA). By replacing Rend with a domain able to multimerize but without mechano-sensitive properties, force through THATCH can remove Rend puncta from the membrane but cannot dissolve the puncta, resulting in Rend droplets in the cytoplasm (Fig. 5d). Note that membrane localization of Rend is still required for the formation of End4p puncta (Fig. S9). Deletion of USH results in constitutive THATCH_ON and the formation of large End4p droplets in the cytoplasm consistent with our model prediction (Fig. 5e). Partial unfolding of Rend (Fig. 5f; Fig. S9) or deletion of Rend (Fig. 5f) both reduced the recruitment of End4p to endocytic sites, thereby impeding End4p from entering the force transmission cycle. In summary, a faulty circuit by Rend and THATCH results in permanent association with the membrane (Fig. 5c) or with F-actin (Fig. 5d-e), or failure to associate with either the membrane or F-actin (Fig. 5f-g).

**Figure 5.**
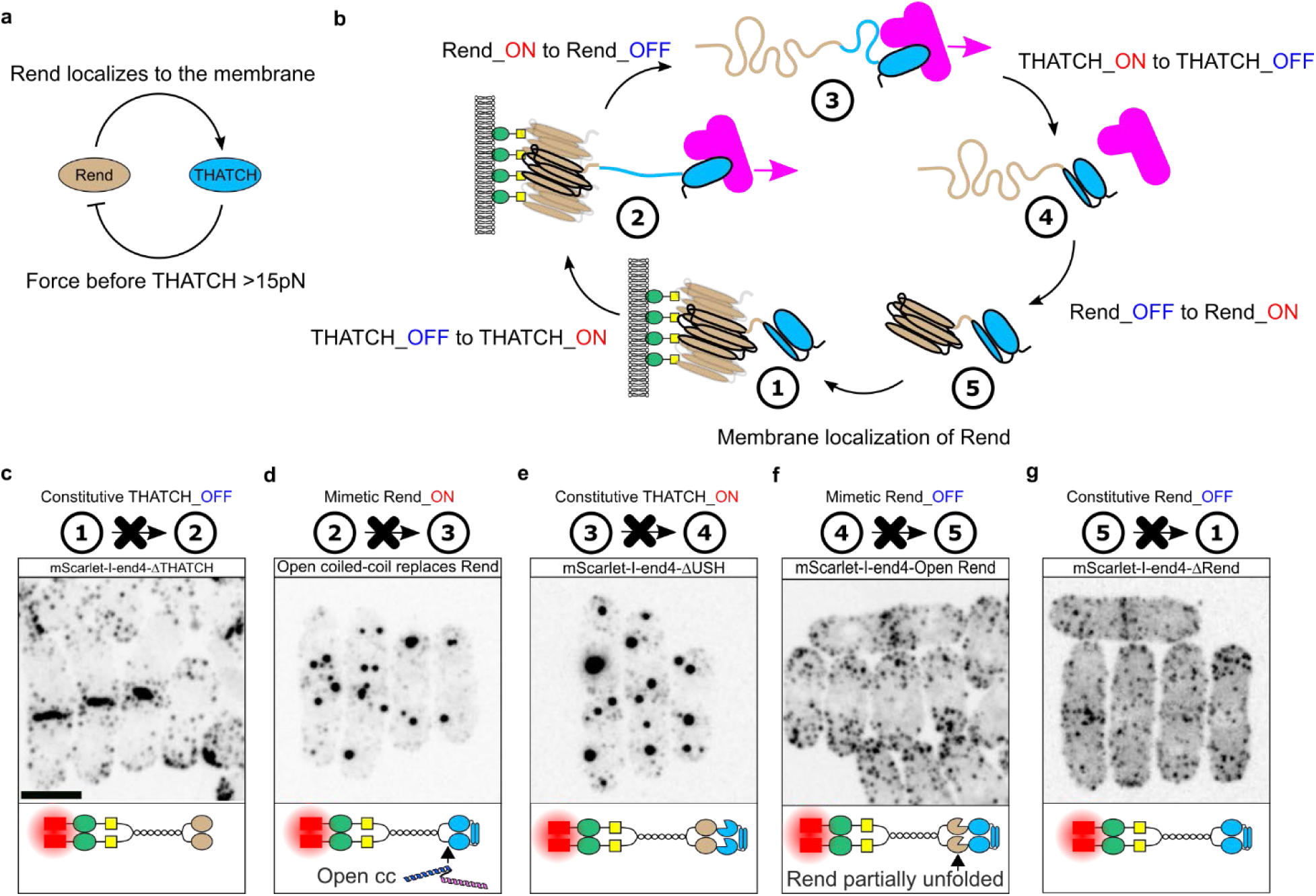
Rend and THATCH forms a mechanical circuit for force transmission. **a**, Mutual regulation of Rend and THATCH forms a circuit. Membrane localization of Rend primes the activation of THATCH, and force through THATCH inactivates Rend at 15pN. **b**, Force transmission cycle by Rend and THATCH in CME. Five states are depicted to indicate the sequential transition of the ON and OFF states of Rend and THATCH. The cycle starts from the transition of state #5 to state #1, and proceeds irreversibly. State #1 (Rend_ON, THATCH_OFF): End4p is localized to the membrane, and Rend forms puncta; State #2 (Rend_ON, THATCH_ON): THATCH is activated and forms stable binding with F-actin. Forces are transmitted through THATCH to Rend and further into the endocytic coat to deform the membrane; State #3 (Rend_OFF, THATCH_ON): Force on Rend exceeds 15pN and Rend unfolds. Tension between Rend and THATCH drops; State #4 (Rend_OFF, THATCH_OFF): Force on THATCH drops to below 8pN. THATCH refolds and unbinds F-actin. Tension between Rend and THATCH continues to drop; State #5 (Rend_ON, THATCH_OFF): Rend refolds and End4p is ready to enter another force transmission cycle. Note that state #2, #3 and #4 are unstable states. End4p domains are simplified and not drawn to scale. **c**, Deletion of THATCH led to the formation of large End4p puncta at the membrane, because THATCH activation is needed for the transition from state #1 to state #2. **d**, Replacing Rend with an open coiled-coil mimics constitutive Rend_ON, where the intermolecular interaction between End4p cannot be terminated by force. This led to the formation of End4p droplets in cytoplasm. **e**, Deletion of USH led to constitutive THATCH_ON, which led to the formation of End4p droplets in cytoplasm. **d** and **e** had similar phenotypes because constitutive Rend_ON leads to THATCH_ON, and state #2 and #3 are unstable states. **f**, Replacing Rend with a partial open version of Rend prevents the refolding of Rend, and led to a higher cytoplasmic ratio of End4p and less End4p at endocytic sites (see also Fig. S?). **g**, Deletion of Rend prevents the formation of Rend puncta at the membrane, and resulted in a higher cytoplasmic ratio of End4p and less End4p at endocytic sites (see also Fig. S4**d**-**e**). **f** and **g** had similar phenotypes because state #4 is an unstable state. Scale bar in **c** applies to **c-g**, 5 μm.

To quantitatively probe the functionality of the Rend-THATCH circuit for force transmission, we developed a mathematical model to determine the response of End4p to varying forces (Fig.6a). We restricted our study to the lifecycle of a fixed pool of membrane bound End4p and actin filaments at an endocytic site. In this model, we considered the Rend domain as folded (R+) or unfolded (R-), and the THATCH domain as bound to an actin filament (T+) or not (T-). Before binding to any actin filament an End4p molecule is in the R+T-state. It captures an actin filament according to mass action kinetics and THATCH immediately becomes catch bond (R+T+), since we assume actin filaments continuously produce force (in other words, End4p acts as a clutch). End4p can detach from actin filaments as a function of force, following a catch bond and slip bond behavior (we used realistic parameters inspired by the literature(*22*, *51*, *52*)). The Rend domain unfolds with a rate following a step function that transitions around 15 pN as measured in this paper (R-T+). At this point End4p does not bear force anymore since it is buffered by the unfolded Rend, and the actin filament is released (R- T-). Rend refolds rapidly (R+T-) and is ready to restart the cycle and bind a new actin filament. The force transmitted to the endocytic membrane can be calculated as the force produced by actin times the ratio of End4p attached to actin filament in a force bearing configuration (i.e., R+T+).

We simulated the model assuming the force produced by the actin filaments is proportional to the amount of actin at endocytic sites, i.e., the input force from F-actin followed a smooth ramp of force from 0 to 20 pN before it decreased (Fig.6b). The transmitted force rapidly plateaued for most of the time course. Both types of bonds (slip bond and catch bond) between THATCH and F-actin were buffered. This result demonstrates that the Rend-THATCH connection buffers the force transmission from F-actin towards the membrane. To better understand the buffering response, we performed simulations using a step function of different magnitude as input force (Fig. 6c). In all cases, the system reached steady state in less than 1 second and the steady-state transmitted force varied non-monotonically with the input force (Fig. 6d). The slip bond scenario led to very inefficient force transmission that became worse with increasing forces. The catch bond scenario allowed force buffering for intermediate forces and sustained force transmission for higher forces, with an efficiency plateauing around 30% for forces above Rend’s unfolding force threshold. Our simulations also showed that Rend unfolding was critical for force buffering since without Rend unfolding (e.g., in RendΔ mutants) the transmitted force keeps increasing with an increase in the input force in the catch bond scenario. Combining catch bond from THATCH and force buffering from Rend allows End4p to dramatically buffer the transmitted force – between 9 and 30 pN of input force (i.e., a ∼3-fold range), the transmitted force is buffered between 5 and 7 pN (i.e., within ∼15% of an average value). Our simulations therefore demonstrate that the Rend-THATCH connection, by combing the catch bond mechanism and mechanical unfolding, filters the stochastic and “noisy” input force from F-actin assembly to a smooth and sustained force transmission towards the membrane.

**Figure 6:**
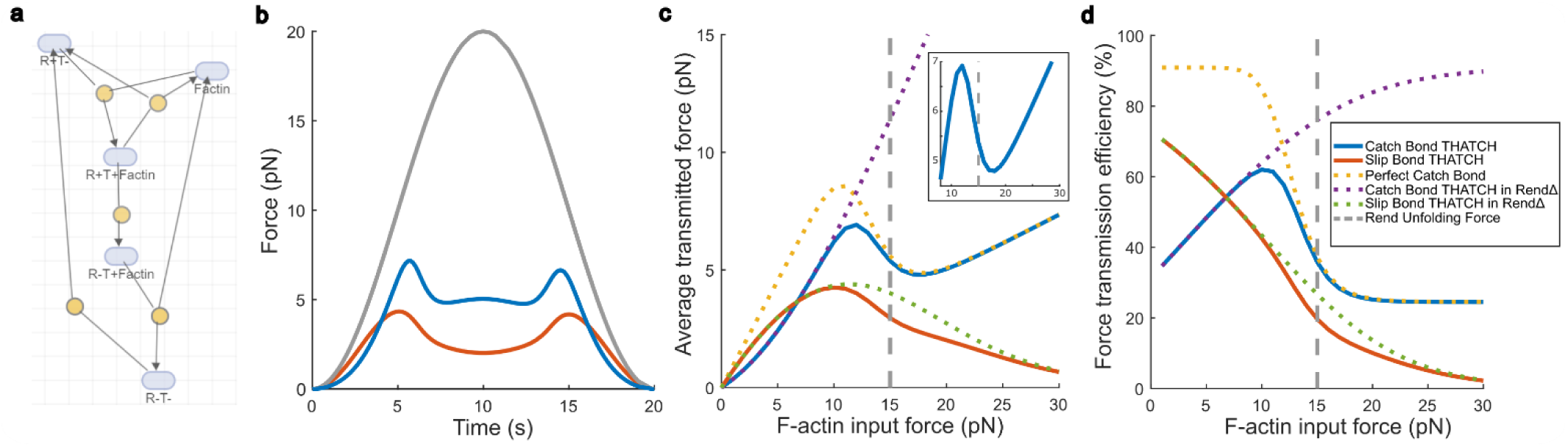
Coupling between THATCH catch bond and Rend mechanosensitive unfolding buffer the forces transmitted to the endocytic machinery. **a**, Schematic of the mathematical model used to simulate the coupling between THATCH and Rend. R+: Folded Rend (Rend_ON); R-: Unfolded Rend (Rend_OFF); T+: catch bond THATCH (THATCH_ON); T-: slip bond THATCH (THATCH_OFF). **b**, Force response out of the Rend-THATCH connection. The input force from F-actin is transmitted from THATCH to Rend and the output force is recorded at the membrane-proximal end of Rend. The input force is roughly proportional to the actin present at endocytic sites. Gray: Input force; Blue: Transmitted force with THATCH as a catch bond; Orange: Transmitted force with THATCH as a slip bond. **c**, Transmitted force by the Rend-THATCH connection as a function of input force produced by actin. The transmitted force is defined as the product of the input force and the ratio between actin-bound THATCH to total actin. Inset: zoom in the region of maximum force buffering (note that the input force varies by ∼100% but the transmitted force varies by ∼15% or less). **d**, Force transmission efficiency as a function of input force. **c-d**, Blue: Catch bond THATCH; Orange: Slip bond THATCH; Yellow: Ideal catch bond that never detaches under force; Purple: Catch bond THATCH in RendΔ mutant; Green: Slip bond THATCH in RendΔ mutant; Gray line: Input force; Vertical dashed line: Rend unfolding force.

## Discussion

The mechanical stabilities of R domains from talin have been characterized in great detail(*23*, *47*, *53–55*), and the formation of catch bond between talin’s THATCH (and other structurally similar five-helix-bundles) and F-actin have been investigated in comparable depth(*18*, *19*, *22*, *25*, *26*). However, the direct connection between R domains and THATCH is key to understanding the initiation, transmission and termination of mechanical forces *in vivo* (Fig. 5b; Fig. 6). In this study, we show that the previously unknown R domain in End4p, Rend, is critical for promoting the binding between THATCH and F-actin to initiate force transmission, for spatially coordinating force transmission, and for terminating force transmission after reaching a hard-wired magnitude. The mechanical circuit formed by Rend and THATCH ensures robust force transmission during CME.

Several factors are crucial for building a functioning circuit using Rend and THATCH. Because forces are transmitted from THATCH to Rend, the mechanical stability of Rend needs to be higher than that of THATCH to allow stable force transmission towards the membrane (Fig. 1d; Fig. 3c). A low mechanical stability of Rend turns THATCH off prematurely. Replacing Rend with R3 from talin, a domain that unfolds at 5pN(*47*, *56*), led to extremely slow cell growth (Fig. S6). This strains also suffers from the inability of R3 to enrich THATCH near the membrane (Fig. 4e), resulting in 40% less End4p molecule at endocytic sites (Fig. 4g). We tried but failed to replace Rend with R12 from talin, a domain that unfolds at an unknown value between 10pN and 20pN(*47*), and forms puncta near membrane (Fig. 4n). We suspect that an overly stable R domain is detrimental to CME because both R and THATCH will be locked in the ON state (Fig. 5d-e), resulting in lethality.

The hysteresis of Rend and the unfolding distance of Rend and THATCH is critical to the irreversibility of the force transmission cycle (Fig. 3c-d). The unfolding of USH (14nm) could be realized during CME, and this change in distance well matches the difference in averaged HIP1R length (10nm) as revealed by FerriTag in electron microscopy (EM), where extended HIP1R molecules are found close to the tip of the endocytic pit, consistent with the idea that force through THATCH is mainly transmitted to the tip region for membrane invagination(*57*). In contrast, EM data of endocytic sites in yeast shows that the Rend may not be fully extended because its unfolding distance (59nm) would make End4p extend beyond the actin meshwork around the endocytic structure(*58*, *59*). The fast refolding of USH prevents the full extension of Rend (Fig. S4). Because Rend unfolds all-or-none and refolds when the force on Rend vanished (<2pN), the drop in tension in the Rend and THATCH connection cannot be reversed before THATCH detaches from F-actin (Fig. 3e). A higher refolding force for USH (8pN) than for Rend (<2pN) therefore prevents the re-engagement of THATCH to F-actin once sufficient force transmission has been achieved (Fig. 5b).

Our *in vivo* force measurement shows that the peak force before THATCH is ∼19pN, which is large enough to be transmitted by a THATCH_ON(*7*). The peak force before Rend is 14-18pN, in agreement with the force buffering of Rend at 15pN (Fig. 3f), and higher than the peak force on End4p close to the clathrin lattice (11pN). These large forces can be well resolved by our coiled-coil force sensors and exceed the range of other *in vivo* force sensors(*60*, *61*). The mechanical stabilities of Rend and THATCH are likely tuned by evolution to match their roles in CME. An interesting prediction is that the mechanical stabilities of Rend and THATCH in Sla2p and HIP1R will be lower, because less mechanical force is needed for CME in budding yeast and in mammalian cells(*4*, *33*, *62*, *63*). However, the unfolding force of Rend is expected to be higher than the unfolding force of THATCH, which will in turn be higher than the refolding force of Rend for the same circuit to function. Recent work by Owen *et. al.* suggests that the unfolding of USH from talin THATCH might be around 5pN(*22*), which makes us speculate that the *in vivo* force before talin THATCH might be 5-10pN based on the estimated mechanical stability of R12.

The ability of Rend to form puncta close to the membrane is intriguing. This property offers a simple mechanism to initiate the binding between THATCH and F-actin by increasing its local concentration and by rerouting force transmission (Fig. 4o). The multiple membrane localization signals in End4p outside of Rend likely regulates the formation of Rend puncta(*34*, *35*, *39*). The spherical droplets formed by dimerized Rend close to the membrane is suggestive of a liquid like material property (Fig. 4i). Cell adhesion proteins including talin and vinculin has been recently shown to undergo membrane-induced 2D phase separation(*64*, *65*), and several endocytic proteins has been found to indeed form condensates to either initiate CME or to cause membrane bending and scission(*66–69*). Non-coding mRNAs have been found in focal adhesions to promote protein condensation(*70*). We have found that several endocytic proteins, including early coat proteins Ede1p, Yap18p, Syp1p and late coat protein End3p, are recruited to the droplets formed by End4p, while late coat proteins Ent1p, Shd1p and Pan1p are not (Fig. S11). How Rend puncta contributes to membrane deformation is an attractive research direction. Condensate or not, the self-association of Rend close to the membrane primes the binding of THATCH to F-actin. This is reminiscent of the transmembrane signaling by other membrane receptors such as integrin, immune receptors, and Receptor tyrosine kinases (RTKs), where clustering is a prerequisite to their activation for downstream signaling(*11*, *71*, *72*). The neighboring THATCH and F-actin binding may be enhanced in a cooperative manner, as has been shown for the αE-catenin actin-binding domain(*19*). We speculate that the cooperativity of Rend domains and that of THATCH domains help accelerate the binding between THATCH and F-actin to achieve switch like engagement between the endocytic coat and the F-actin meshwork. Moreover, a layer of self-associating Rend spatially coordinates the transmission of forces towards the endocytic coat and filters the F-actin fluctuations in the outmost layer (Fig. 6). Additional layers made by protein-protein interactions likely exists in the endocytic coat, of which the clathrin lattice is an obvious candidate. Forces transmitted from F-actin to the endocytic coat are thus collected and redistributed through several layers before reaching the membrane(*7*).

The unfolding of Rend by mechanical forces retrieves Rend from the membrane, and the putative phosphorylation of Rend by Ark/Prk kinases at T841 may be involved in dissolving Rend puncta during CME, although mechanical forces seem to play a major role(*48*, *73*) (Fig. S7). The regulation of Rend puncta during CME by both mechanical and chemical signals is an interesting direction for future exploration, and there may be mutual regulations between mechanical forces, protein crowding and kinase specificity/activity(*74*, *75*). The unfolding of Rend causes a localized mechanical disconnection between the endocytic coat and the F-actin right at the tip of the endocytic pit, which could be used to release the energy stored in cross-linked F-actin to pinch off the endocytic vesicle after sufficient energy accumulation(*58*, *76*). Quantitative modeling of this process, one which adds spatial dimensions to what has been modeled in this study (Fig. 6), will be informative in explaining the later stages of CME.

The discovery of Rend on End4p imply an evolutionary connection between End4p/Sla2/HIP1R and talin. Both are linear molecules that bind the membrane at the N-terminus, contains R domain(s) in the middle, and bind to F-actin at the C-terminus through THATCH (Fig. S12). We show also that unfolded Rend, but not the folded Rend, binds to vinculin head domain 1 (Vd1) in fission yeast cells (Fig. S13). Unknown binding partners of Rend before and after unfolding may help regulate CME, analogous to force-dependent protein binding/unbinding to talin R domains(*27*, *53*). Because endocytosis likely predates multicellularity, cells may have co-opted the endocytic machinery to build cell-matrix or cell-cell connections during evolution. Consistent with this idea, in a recent lab evolution experiment, several genes related to endocytosis and cytoskeletal organization are implicated in the transition into multicellular life cycle(*77*). One fungus species, *Allomyces macrogynus*, contains both HIP1R and talin, and structural predictions suggest they each contain 2 and 13 R domains (including THATCH)(*78*). It will be informative to explore if R domain duplication or deletion led to the apparent functional homology between End4p/Sla2/HIP1R and talin. It will also be meaningful to check if similar mechanical circuits represent a general solution for robust mechanotransduction *in vivo*.

## Notes

### Competing Interest Statement

The authors have declared no competing interest.

